# Extensive CD8β depletion does not prevent control of viral replication or protection from challenge in macaques chronically infected with a live attenuated simian immunodeficiency virus

**DOI:** 10.1101/608554

**Authors:** Matthew S. Sutton, Amy Ellis-Connell, Alexis J. Balgeman, Gabrielle Barry, Andrea M. Weiler, Scott J. Hetzel, Yan Zhou, Annie Kilby, Rosemarie D. Mason, Kristin K. Biris, John R. Mascola, Nancy J. Sullivan, Mario Roederer, Thomas C. Friedrich, Shelby L. O’Connor

**Affiliations:** Department of Pathology and Laboratory Medicine, UW-Madison, Madison, WI; Wisconsin National Primate Research Center, UW-Madison, Madison, WI; Department of Biostatistics and Medical Informatics, UW-Madison, Madison, WI; Department of Pathobiological Sciences, UW-Madison, Madison, WI; Vaccine Research Center, National Institute of Allergy and Infectious Diseases, National Institutes of Health, Bethesda, MD

## Abstract

We evaluated the contribution of CD8αβ+ T cells on control of live-attenuated simian immunodeficiency virus (LASIV) replication during chronic infection and subsequent protection from pathogenic SIV challenge. Unlike previous reports with a CD8α-specific depleting monoclonal antibody (mAb), the CD8β-specific mAb CD8β255R1 selectively depleted CD8αβ+ T cells without also depleting non-CD8+ T cell populations that express CD8α, such as natural killer (NK) cells and γδ T cells. Following infusion with CD8β255R1, plasma viremia transiently increased coincident with declining peripheral CD8αβ+ T cells. Interestingly, plasma viremia returned to pre-depletion levels even when peripheral CD8αβ+ T cells did not. Although depletion of CD8αβ+ T cells in the lymph node (LN) was incomplete, frequencies of these cells were three-fold lower (p=0.006) in animals that received CD8β255R1 compared to control IgG. It is possible that these residual SIV-specific CD8αβ+ T cells may have contributed to suppression of viremia during chronic infection. We also determined whether infusion of CD8β255R1 in the LASIV-vaccinated animals increased their susceptibility to infection following intravenous challenge with pathogenic SIVmac239. We found that 7/8 animals infused with CD8β255R1, and 3/4 animals infused with the control IgG, were resistant to SIVmac239 infection. These results suggest that infusion with CD8β255R1 did not eliminate the protection afforded to LASIV vaccination. This provides a comprehensive description of the impact of CD8β255R1 infusion on the immunological composition of the host, when compared to an isotype matched control IgG, while showing that the control of LASIV viremia and protection from challenge can occur even after CD8β255R1 administration.

**Importance:** Studies of SIV-infected macaques that deplete CD8+ T cells *in vivo* with monoclonal antibodies have provided compelling evidence for their direct antiviral role. These studies utilized CD8α-specific mAbs that target both the major (CD8αβ+) and minor (CD8αα+) populations of CD8+ T cells, but additionally deplete non-CD8+ T cell populations that express CD8α, such as NK cells and γδ T cells. In the current study, we administered the CD8β-specific depleting mAb CD8β255R1 to cynomolgus macaques chronically infected with a LASIV to selectively deplete CD8αβ+ T cells without removing CD8αα+ lymphocytes. We evaluated the impact on control of virus replication and protection from pathogenic SIVmac239 challenge. These results underscore the utility of CD8β255R1 for studying the direct contribution of CD8αβ+ T cells in various disease states.

## Introduction

Multiple lines of evidence suggest that CD8+ T cells contribute to control of virus replication and subsequently influence disease progression following human immunodeficiency virus (HIV) infection. For instance, the emergence of HIV-specific CD8+ cytotoxic T lymphocytes (CTLs) during acute infection is temporally associated with decreases in peak viral load and the decline of viral replication to set point viral load (1, 2). Indeed, CD8+ CTLs have been shown to lyse HIV-infected cells *in vitro*, and the selective pressure they exert *in vivo* often leads to the emergence of immune escape variants (3–7). The strongest argument comes from studies of macaques infected with simian immunodeficiency virus (SIV) that are infused with a monoclonal antibody (mAb) that is specific for the CD8α molecule of CD8+ lymphocytes. Following infusion with this antibody, depletion of CD8α+ cells persists for approximately 2 to 4 weeks and is accompanied by a transient increase in virus replication until control is regained coincident with the reemergence of CD8α+ lymphocytes (8–12, 14, 31, 45–48, 53–54). Of note, control of virus replication is lost following *in-vivo* depletion of CD8+ lymphocytes even during antiretroviral therapy (ART), further suggesting that functional CD8+ T cells are needed to maintain effective viral control even while on ART (11, 12). Notably, however, CD8α-specific mAbs deplete not only CD8+ T cells, but also a variety of cell populations that express the CD8α molecule.

The CD8 molecule is expressed as either a CD8αα homodimer or a CD8αβ heterodimer on the cell surface and is present on lymphocytes of both the innate and adaptive immune system (15, 16, 18, 49). The most common lymphocytes to express CD8 are conventional CD8+ T cells (TCRαβ+CD3+), which can be divided into a major population that express CD8αβ and a minor population that express CD8αα (13). There also exist populations of TCRγδ+CD3+ T cells and CD3-natural killer (NK) cells that express CD8αα (17, 18, 52). γδ T cells (TCRγδ+CD3+CD8αα+) which comprise ∼ 6% of CD3+ T cells (17) can block HIV-1 entry via the secretion of β-chemokines (23), enhance antibody-dependent cellular cytotoxicity (ADCC) (24), and directly lyse HIV-infected cells (25). NK cells (CD3-CD8αα+) comprise ∼16% of peripheral lymphocytes, and have recently been reported to possess traits of adaptive immunity that may contribute to control of HIV-1 replication (19, 20). Accordingly, the contribution of conventional CD8+ T cells to viral control are complicated by the depletion of additional cell populations that express CD8α when using CD8α-depleting mAbs (10, 14, 53). One approach to better define the antiviral role of CD8+ T cells *in vivo* is to administer a CD8β-specific depleting mAb, as this should selectively deplete CD8αβ+ T cells without removing CD8αα+ lymphocytes or other non-T cell populations. Indeed, two recent studies using the CD8β-specific mAb CD8β255R1 in rhesus macaques provide evidence that CD8αβ+ T cells can be specifically depleted *in vivo* (40, 41).

Macaques vaccinated with SIVmac239Δnef, a live-attenuated SIV (LASIV) variant of pathogenic SIVmac239, are useful for evaluating the role of CD8αβ+ T cells in control of virus replication and protection from SIV challenge. Although rare hosts spontaneously control pathogenic HIV or SIV in a manner dependent on particular major histocompatibility complex (MHC) alleles, control of SIVmac239Δnef replication occurs in nearly every vaccinated animal, regardless of host MHC genetics (27–31). These observations question whether the contribution of conventional CD8αβ+ T cells to control of SIVmac239Δnef is equivalent to their contribution to control of pathogenic SIV. Moreover, vaccination with SIVmac239Δnef is the most successful example of vaccine-induced protection from challenge with homologous SIV strains and, less frequently, from challenge with heterologous SIV strains (31–34). After more than 25 years of effort, the precise immune mechanism(s) responsible for this protection are still under debate (35– 39). Thus, defining whether SIVmac239Δnef-mediated vaccine protection requires CD8αβ+ T cells may also help inform either therapeutic or prophylactic HIV vaccine design.

In this study, we utilize the CD8β-specific mAb CD8β255R1 to specifically deplete CD8αβ+ T cells in LASIV-vaccinated Mauritian cynomolgus macaques (MCMs) (42), and then measure the impact on control of LASIV replication and protection from pathogenic SIV challenge. In contrast to two recent studies that evaluated the impact of CD8β255R1 in SIV-infected or SHIV-infected rhesus macaques (40, 41), we include animals treated with a control IgG to distinguish those immunological effects specific to CD8β-depletion from those that are a result of infusion of a nonspecific IgG antibody. We also determine whether depletion of CD8αβ+ T cells affects the frequency and proliferative capacity of CD8αα+ T cells, NK cells, and γδ T cells. Thus, we expand the current knowledge of the immunological effects that follow infusion with CD8β255R1 and provide the first comprehensive comparison to an isotype matched control IgG.

## Materials and Methods

### Animal care and use

Two Vietnamese-origin cynomolgus macaques were purchased from Covance Inc. and included in the depletion study of naïve cynomolgus macaques. Both animals were housed and cared for at NIHAC-Poolesville (National Institutes of Health Animal Center) according to protocols approved by the Vaccine Research Center Animal Care and Use Committee. Both animals received a single 50mg/kg intravenous infusion of the anti-CD8β monoclonal antibody (mAb) CD8β255R1.

Eleven Mauritian cynomolgus macaques (MCMs) were purchased from Bioculture Ltd. and included in the depletion/challenge study of cynomolgus macaques chronically infected with a LASIV. These MCMs were housed and cared for by the Wisconsin National Primate Research Center (WNPRC) according to protocols approved by the University of Wisconsin Graduate School Animal Care and Use Committee. All eleven animals were previously infected intravenously with 10ng p27 of wild-type SIVmac239Δnef (cy0749 and cy0752) or a variant of SIVmac239Δnef containing 10 nonsynonymous mutations (cy0685, cy0688, cy0690, cy0691, cy0750, cy0753, cy0755, cy0756, and cy0757) for 34 to 73 weeks prior to the start of this study (42). All animals were homozygous for the M3 MHC haplotype, with the exception of cy0691 that was heterozygous for the M2 and M3 MHC haplotypes. Seven MCMs received a single 50mg/kg intravenous infusion of the anti-CD8β monoclonal antibody (mAb) CD8β255R1 and four MCMs received a single 50mg/kg intravenous infusion of the DSPR1 rhesus recombinant IgG control mAb, both of which were provided by the NIH Nonhuman Primate Reagent Resource (R24 OD010976, U24 AI126683). All eleven MCMs were challenged intravenously with 100TCID50 SIVmac239 (0.71ng p27) four weeks following mAb infusion.

### Antibodies

CD3-AF700 (clone: SP34-2; BD Biosciences), CD4-BV711 (clone: OKT4; BioLegend), CD8α-PE (clone: DK25; Dako), CD8β-ECD (clone: 2ST8-5H7; Beckman Coulter), TCRγδ-FITC (clone: 5A6.E9; Invitrogen), CD95-PE/Cy5 (clone: DX2; BD Biosciences), CD28-BV510 (clone: CD28.2; BD Biosciences), CCR7-Pacific Blue (clone: G043H7; BioLegend), NKG2a-PE/Cy7 (clone: Z199; Beckman Coulter), CD16-BV650 (clone: 3G8; BD Biosciences), CD16-BV786 (clone: 3G8; BD Biosciences), and Ki-67-AF647 (clone: B56; BD Biosciences). LIVE/DEAD Fixable Near-IR Dead Cell Stain (Invitrogen) was used to assess cell viability. All samples were run on the LSR II (BD Biosciences) and analyzed using FlowJo, Version 9.9.6 (BD Biosciences). The presence of the CD8β255R1 mAb does not cross-block binding with the anti-CD8β clone 2ST8-5H7 (40).

### Peptides

The NIH AIDS Research and Reference Reagent Program (Germantown, MD) provided 15-mer peptides overlapping by 11 amino acid positions that spanned the entire SIVmac239 proteome. Gag and Env peptides were combined into 2 and 3 pools, respectively, and used at a final pooled concentration of 5μM during stimulation.

### Lymphocyte isolation and phenotype staining

In the depletion study of naïve cynomolgus macaques, phenotype staining of peripheral blood mononuclear cells (PBMCs) was performed in triplicate on whole blood and in duplicate on lymph node samples. Cells were incubated with a surface antibody mix for 25 minutes in the dark at room temperature. In the depletion/challenge study, PBMC were isolated from EDTA-anticoagulated blood by ficoll-based density centrifugation as previously described (34). Cells were resuspended in RPMI 1640 (HyClone) supplemented with 10% fetal calf serum (FCS), 1% antibiotic-antimycotic (HyClone), and 1% L-gluatmine (HyClone) (R10 medium). Approximately 0.5-1.0 x 10^6^ PBMC were placed in cluster tubes (Fisher Scientific) and washed with 1x phosphate-buffered saline (PBS) prior to staining. Surface antibodies were added and cells were incubated for 30 min in the dark at room temperature. Cells were then washed twice with 1x PBS supplemented with 10% FCS (10% FCS/PBS), and fixed with 2% paraformaldehyde (PFA) for a minimum of 30 minutes. Fixed cells were then washed once with 1x PBS and then permeabilized with Medium B (Invitrogen, Carlsbad CA) and allowed to incubate with intracellular antibodies for 30 min in the dark at room temperature. Cells were then washed twice with 10% FCS/PBS and resuspended in 2% PFA prior to data collection. Following exclusion of doublets and dead cells, lymphocyte populations were defined as follows: CD8αβ+ T cells: CD3+CD4-TCRγδ-CD8α+CD8β+; CD8αα+ T cells: CD3+CD4-TCRγδ-CD8α+CD8β-; γδ T cells: CD3+CD4-CD8β-CD8α+TCRγδ+; NK cells: CD3-CD4-CD8α+NKG2a+CD16+; CD4 T cells (naive): CD3+CD4+CD8α-CD95-CD28+CCR7+; CD4 T cells (effector memory): CD3+CD4+CD8α-CD95+CD28-CCR7-; CD4 T cells (central memory): CD3+CD4+CD8α-CD95+CD28+CCR7+.

### Intracellular cytokine staining (ICS)

Flow cytometry was used to measure intracellular cytokine expression as previously described (59). Cryopreserved cells isolated from LNs were thawed and washed twice in warm R10 medium and allowed to rest at 37°C overnight prior to stimulation. Ki-67 was used to measure the proliferative capacity of lymphocytes freshly-isolated from blood, and IFN-γ, tumor necrosis factor alpha (TNF-α), and CD107a was used to measure SIV-specific responses in cryopreserved lymphocytes isolated from LNs at week 3 post mAb infusion. LN cells were incubated with CD107a and stimulated with Gag or Env peptide pools for a total of 6 hours, with brefeldin A (Sigma-Aldrich) and monensin (BioLegend) added 1 hour after stimulation. PBMC and LN cells were incubated with LIVE/DEAD fixable near-infrared dead cell stain for 20 minutes, incubated with surface markers for 30 minutes, and fixed with 2% paraformaldehyde (PFA) for at least 20 minutes. Medium B (Invitrogen) was used to permeabilize PBMC and 0.1% saponin was used to permeabilize LN cells. PBMC and LN cells were stained for 30 minutes with Ki-67 or IFN-γ and TNF-*α*, respectively. Data were collected on an LSR II instrument (BD Biosciences) using 2% PFA-fixed cells and then analyzed using FlowJo version 9.9.6 (BD Biosciences).

### Plasma viral load analysis

Plasma was isolated alongside PBMC and cryopreserved at −80°C prior to analysis. SIV *gag* viral loads were determined as previously described (34). Briefly, viral RNA (vRNA) was isolated from plasma, reverse transcribed, and amplified with the Superscript III platinum one-step quantitative reverse-transcription-PCR (RT-PCR) system (Invitrogen). The detection limit of the assay was 100 vRNA copy equivalents per mL of plasma (copies/mL). When the viral load was at or below the limit of detection, the detection limit value of 100 was reported. Full length *nef* and Δ*nef* viral loads were determined as previously described (34). Briefly, highly-specific real-time RT-PCR assays were used with primers that accurately differentiate viruses containing full-length *nef* from those that contain *nef* with a 182 base-pair (bp) deletion, using the methods described above. Serial dilutions of *in vitro* transcripts for both full-length *nef* and *nef* with a 182-bp deletion were used as internal standards for each run. The same machines and software used for the *gag* viral load assay were used to detect and quantify the *nef* and Δ*nef* viral loads. The limit of detection was identical to that for the SIV *gag* viral load assay.

### Virus Neutralization Assays

SIV Env pseudoviruses were produced as previously described (43). Briefly, plasmid DNA encoding SIV gp160 was combined with a luciferase reporter plasmid containing the essential HIV structural genes to produce pseudoviruses expressing SIVmac251.H9, SIVmac251.30, SIVsmE660.CP3C, or SIVmac239 Env. Using TZM-bl target cells, virus neutralization was measured following incubation with SIV-Env pseudovirus and plasma collected from blood. The 50% inhibitory dilution (ID50) was defined as the plasma dilution that caused a 50% reduction in relative light units (RLU) compared to virus control wells following subtraction of background RLU. A nonlinear-regression 5-parameter Hill slope equation was used to calculate plasma ID50 values.

### Statistical Analyses

Comparison of two groups in Figures 2b-c, 4c-d, and 5d-f was conducted using Wilcoxon rank sum tests. Comparison of changes from baseline separately for each treatment group shown in Figures 2a, 3, 4a-b, and 5a-c utilized repeated measures ANOVA (RM-ANOVA) to estimate the mean contrasts with animal as a random effect. Comparison between both groups in changes from pre-treatment levels at multiple follow-up time points also utilized RM-ANOVA with treatment, time, and their interaction as fixed effects, and animal as a random effect. Model assumptions of RM-ANOVA were examined and were not deemed to be grossly violated. Dunnett’s p-value adjustment for multiple testing over multiple time points was used to keep a family-wise 0.05 type 1 error rate (58). All tests were conducted using a two-sided significance level of 0.05. All analyses were conducted using R for statistical computing version 3.3 (44).

**Figure 1.**
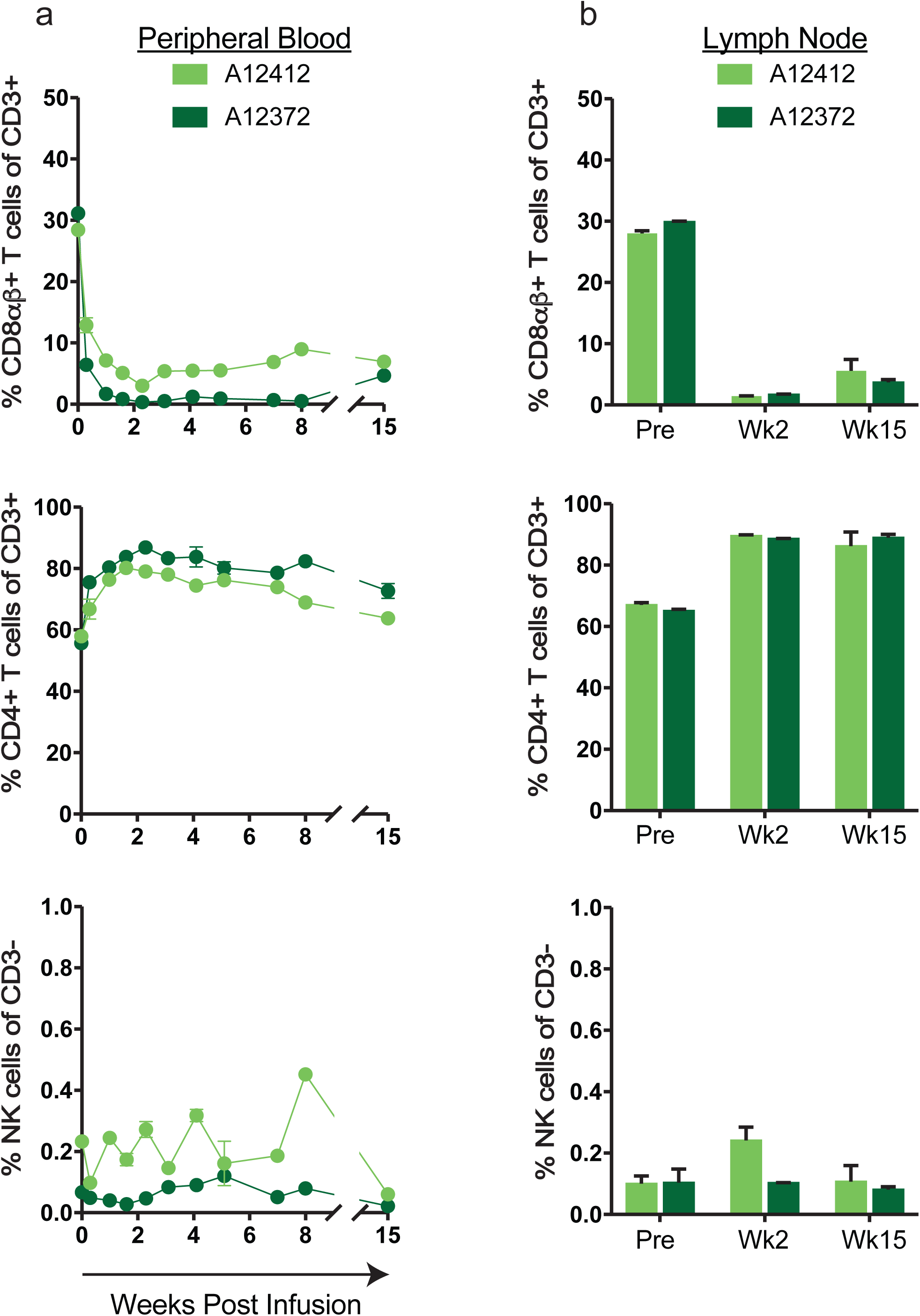
Infusion with CD8β255R1 selectively depletes CD8αβ+ T cells in naïve cynomolgus macaques. Two naïve cynomolgus macaques (A12412 and A12372) received a single 50 mg/kg intravenous infusion with CD8β255R1. Percent CD8β+ T cells (top), CD4+ T cells (middle), and CD16+ NK cells (bottom) in (a) peripheral blood and (b) lymph nodes were measured by multicolor flow cytometry. Data are shown as mean and standard deviation. Peripheral blood samples from each animal were analyzed in triplicate, while lymph node samples from each animal were analyzed in duplicate.

**Figure 2.**
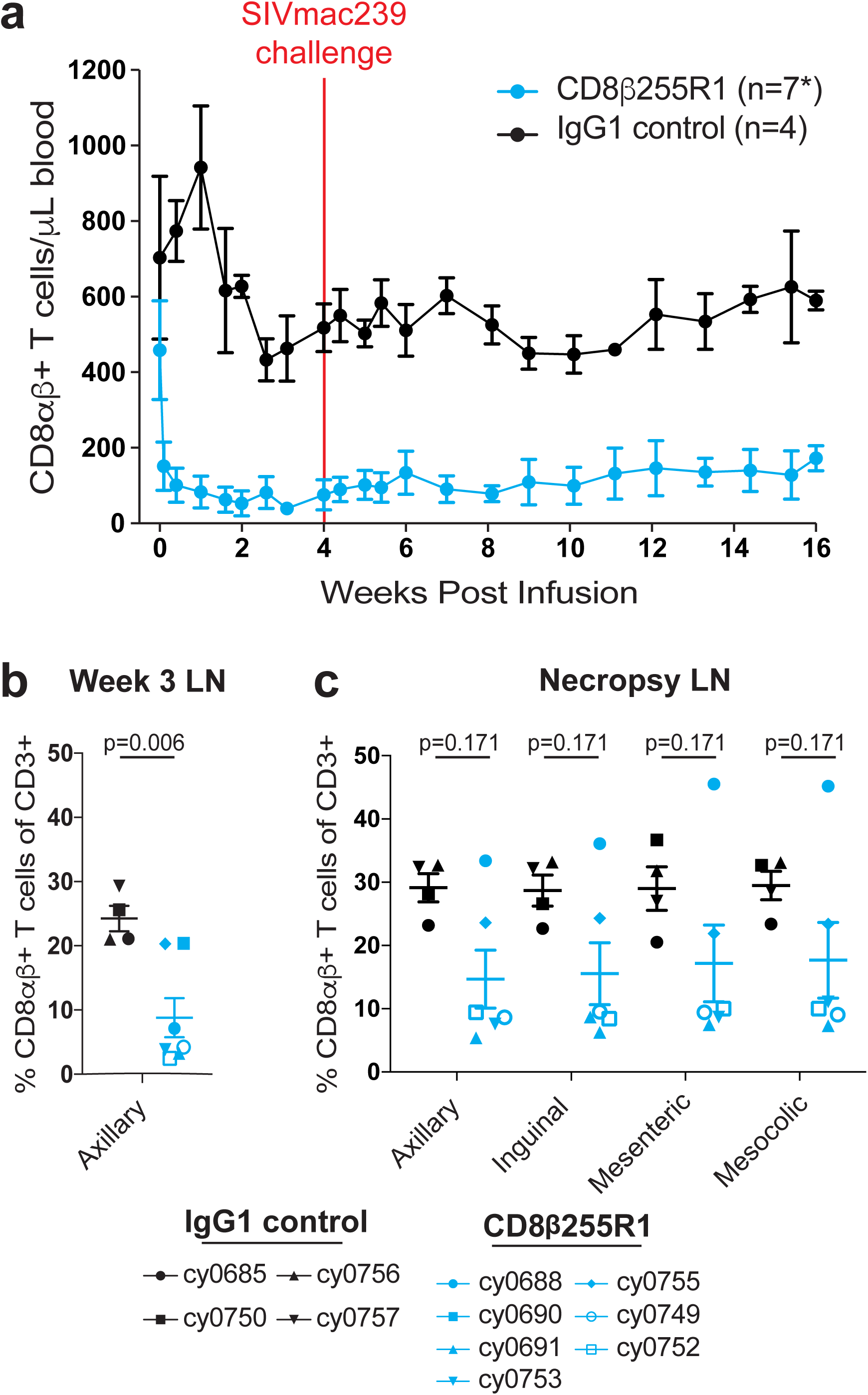
Depletion of CD8αβ+ T cells during chronic LASIV infection. (a) Absolute count of CD8αβ+ T cells were assessed in peripheral blood for 16 weeks following infusion with CD8β255R1 (blue) or control IgG (black). Percent CD8αβ+ T cells from lymph node biopsies at (b) 3 weeks post-infusion and (c) at necropsy (week 17-18). Data are represented as mean and standard error of the mean. *Data shown include four IgG-treated animals and seven CD8β255R1-treated animals through week 6, at which point six CD8β255R1-treated animals are shown for the remainder of the study due to one animal (cy0690) requiring early euthanasia.

**Figure 3.**
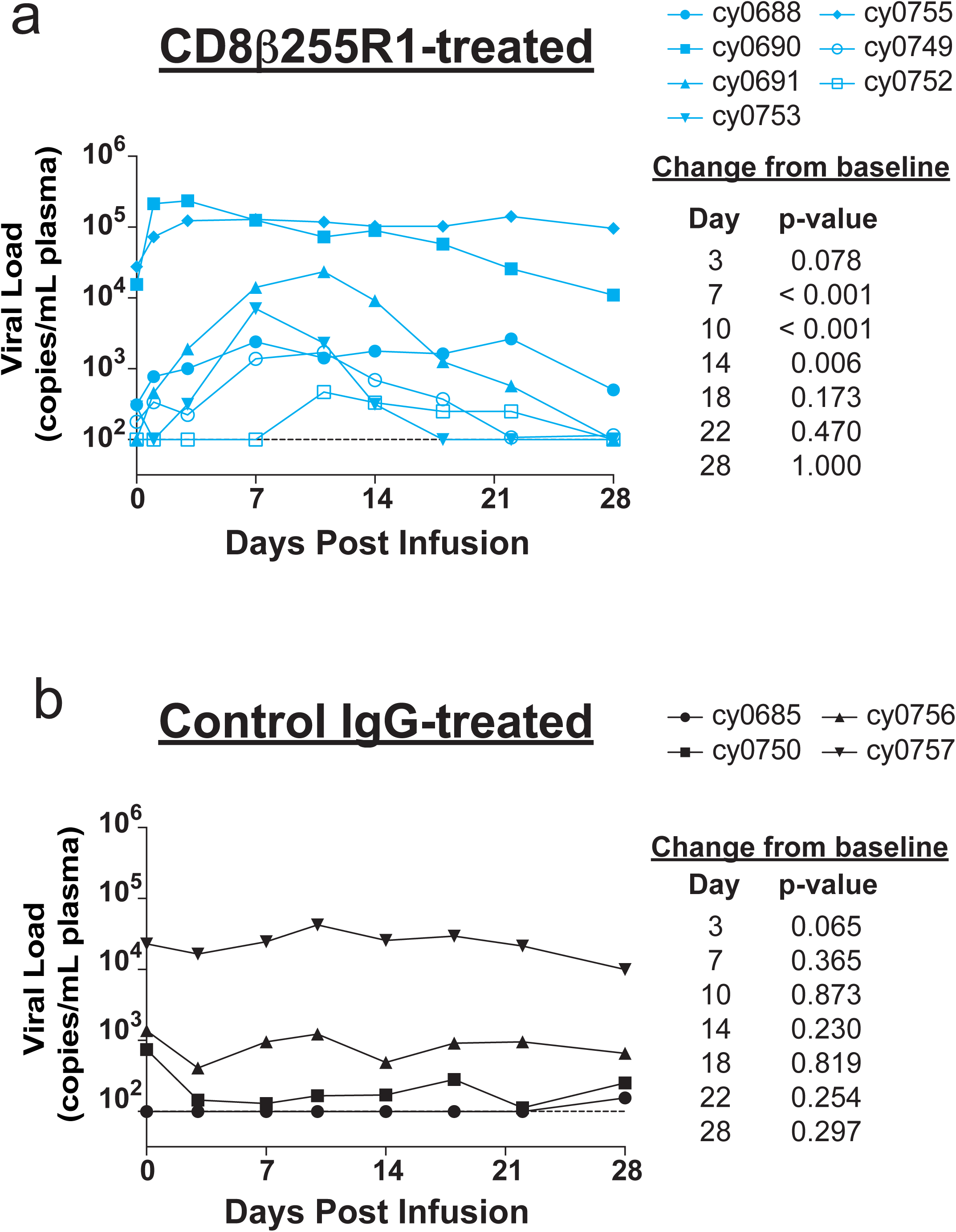
Regained control of viral recrudescence following depletion of peripheral CD8αβ+ T cells. Plasma viral load was measured for four weeks after mAb infusion. (a) Viral load initially increased following infusion with CD8β255R1, but returned to pre-depletion levels by day 18. (b) Viral load did not change significantly in animals that received control IgG. The limit of detection of the plasma viral load assay (100 vRNA copies/mL) is shown with a horizontal dashed line.

**Figure 4.**
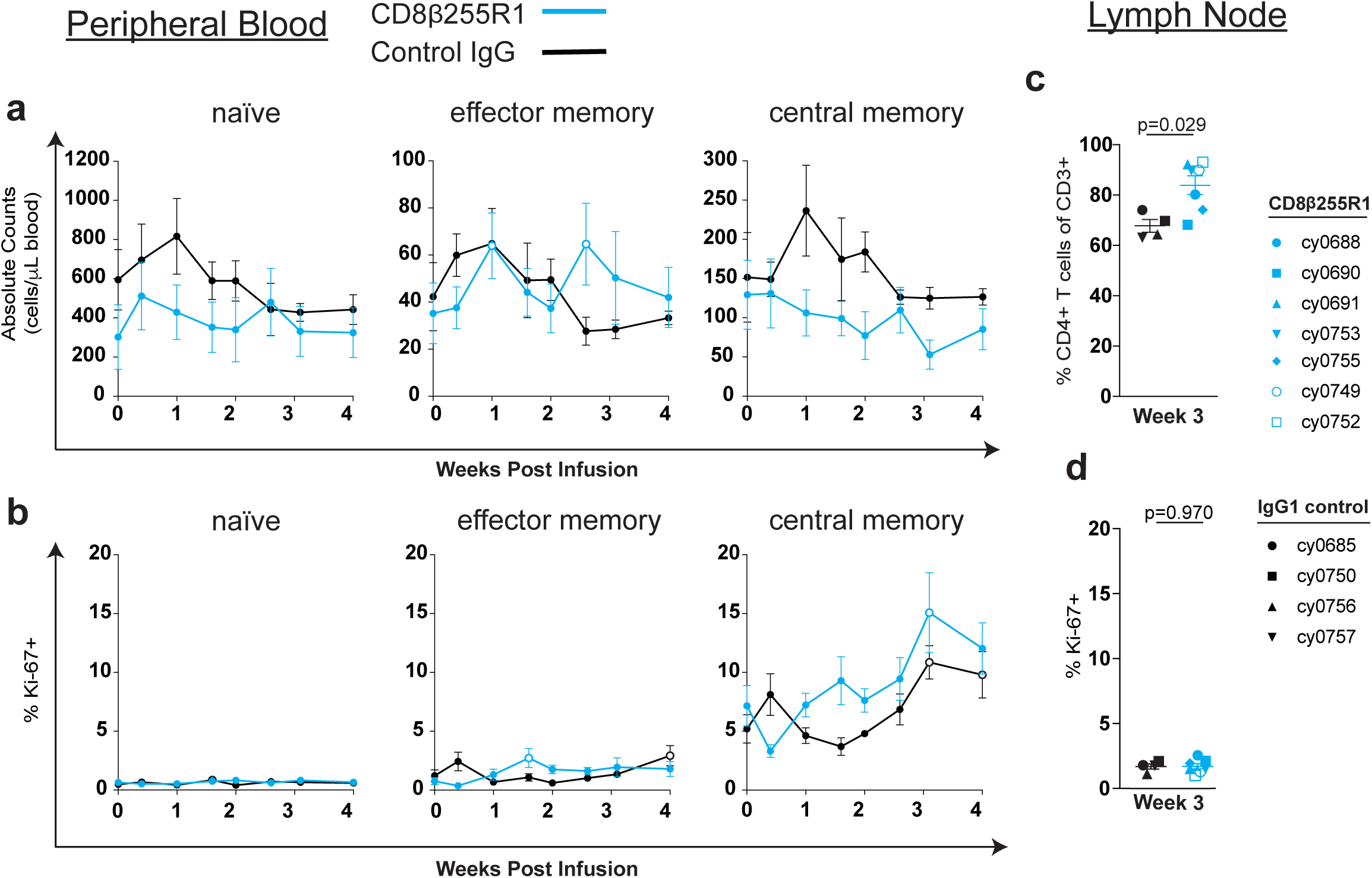
Minimal changes in the frequency and proliferation of CD4+ T cell populations following administration of CD8β255R1. Absolute cell count and percent Ki-67+ of CD4+ T cells from peripheral blood and lymph nodes were measured by multicolor flow cytometry for 4 weeks following mAb infusion. Within peripheral blood, individual time points represent mean ± SEM for animals that received CD8β255R1 (blue) or control IgG (black) and open circles represent a significant (p<0.05) change from baseline. (a) Absolute cell count and (b) percent Ki-67+ of naive (left), effector memory (middle), and central memory (right) CD4+ T cells in peripheral blood. Within the lymph nodes, individual animals are represented by a unique symbol and red text represents a significant difference between groups. (c) Percent CD4+ T cells of CD3+ T cells and (d) percent Ki-67+ of CD4+ T cells from axillary lymph nodes biopsied at week 3 post mAb infusion.

**Figure 5.**
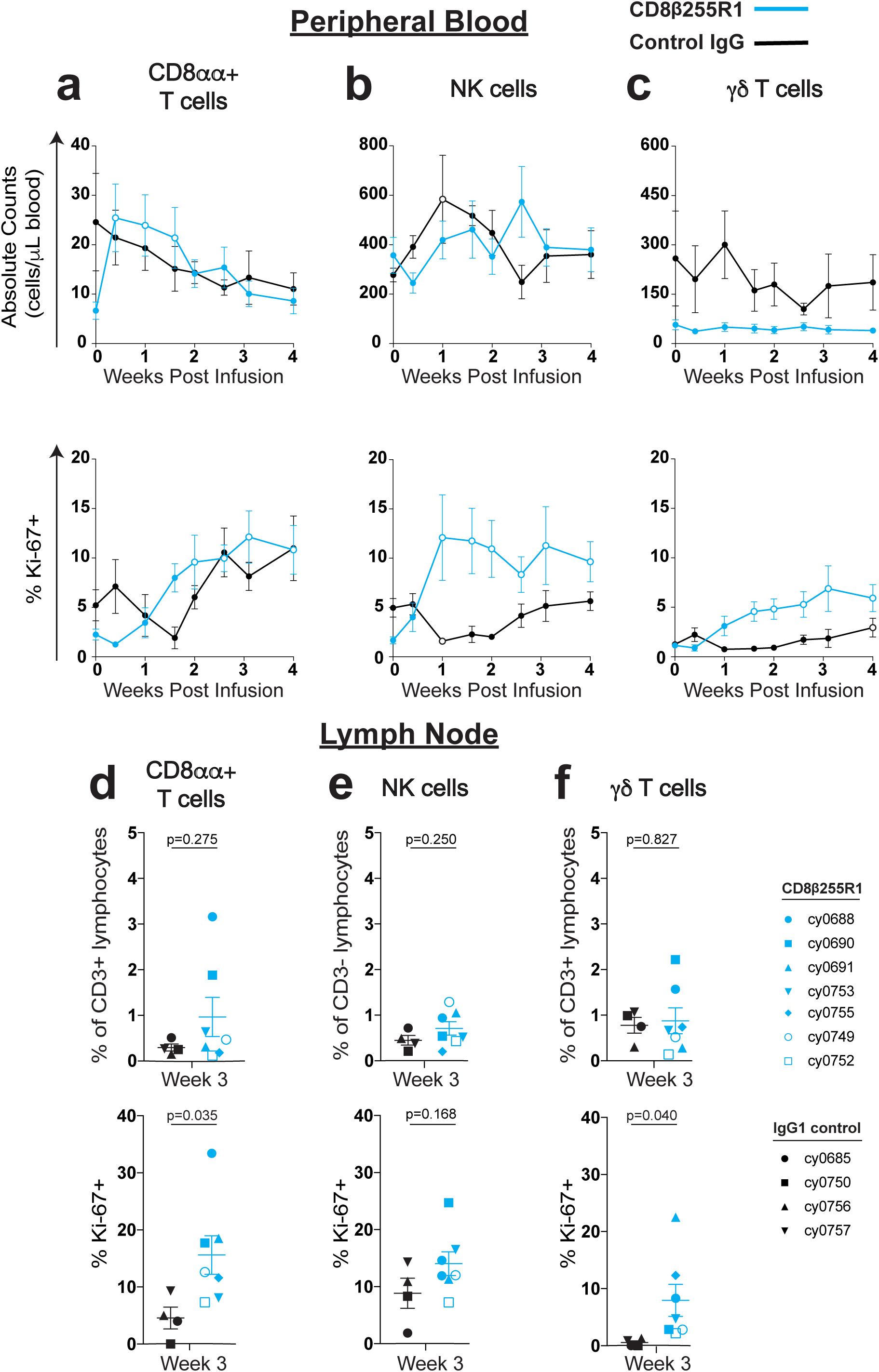
Proliferation of CD8αα+, NK cells, and γδ T cells in peripheral blood following infusion with CD8β255R1. Absolute cell count and percent Ki-67+ of CD8α+ lymphocytes from peripheral blood and lymph nodes were measured by multicolor flow cytometry for 4 weeks after infusion with CD8β255R1 (blue) or control IgG (black). Within peripheral blood, individual time points represent mean ± SEM for animals and open circles represent a significant change from baseline. Absolute count (top) and percent Ki-67+ (bottom) of (a) CD8αα+ T cells, (b) NK cells, and (c) γδ T cells in peripheral blood. Within the lymph nodes, individual animals are represented by a unique symbol and red text represents a significant difference between groups. (d) Percent CD8αα+ T cells of CD3+ lymphocytes (top) and percent Ki-67+ CD8αα+ T cells (bottom) in lymph nodes. (e) Percent NK cells of CD3-lymphocytes (top) and percent Ki-67+ NK cells (bottom) in lymph nodes. (f) Percent γδ T cells of CD3+ lymphocytes (top) and percent Ki-67+ γδ T cells (bottom) in lymph nodes.

## Results

### Infusion with CD8β255R1 selectively depletes CD8αβ+ T cells in naïve cynomolgus macaques

Previous reports of CD8β-depletion following infusion with the anti-CD8β mAb CD8β255R1 were performed exclusively with rhesus macaques (40, 41). To determine whether CD8β255R1 can similarly deplete CD8αβ+ T cells in cynomolgus macaques, two naïve animals (A12412 and A12372) were intravenously infused with a single 50 mg/kg dose of the anti-CD8β mAb CD8β255R1 and evaluated for 15 weeks. Flow cytometry was used to monitor CD8αβ+ T cells in peripheral blood mononuclear cells (PBMC) and peripheral lymph nodes (LNs) before and after infusion in both animals. Depletion of peripheral CD8αβ+ T cells was rapid and sustained (Figure 1a, top) with declines in the percentage of lymphocytes relative to baseline reaching its nadir at week 2 (A12412: 89%, A12372: 99%), and remaining substantially reduced at week 15 (A12412: 75%, A12372: 85%). Levels of circulating CD4+ T cells increased transiently following infusion (Figure 1a, middle), while circulating NK cells were relatively unchanged over the course of 15 weeks (Figure 1a, bottom).

Depletion of LN CD8αβ+ T cells after administration of the CD8β255R1 mAb has been reported in rhesus macaques (http://www.nhpreagents.org/NHP/availablereagents.aspx?Cat=2). We also observed reductions in LN CD8αβ+ T cells (Figure 1b, top) in both cynomolgus macaques at week 2 (mean: 94.5%; range: 94% to 95%) that remained reduced by approximately 84% at week 15 (range: 80% to 88%) when compared to pre-depletion levels. Similar to our observations in the blood, we detected elevated levels of CD4+ T cells in LNs (Figure 1b, middle) that persisted for 15 weeks following infusion while levels of NK cells in LNs (Figure 1b, bottom) remained largely unaffected for 15 weeks. These results are not surprising, as a decrease in the frequency of CD3+ T cells that are CD8+ will correspond to an increase in the frequency of CD3+ cells that are CD4+ in the same compartment. To our knowledge, this is the first assessment of the immunological effects following infusion with CD8β255R1 in cynomolgus macaques, and these results are similar to those observed in rhesus macaques (40, 41).

### Impact of CD8β255R1 infusion on CD8αβ+ T cells during chronic LASIV infection

Seven MCMs chronically infected with either SIVmac239Δnef (n=2) or a SIVmac239Δnef variant (n=5) (42) received a single 50 mg/kg intravenous infusion of the anti-CD8β mAb CD8β255R1. Four MCMs infected with the same SIVmac239Δnef variant received a single 50 mg/kg intravenous infusion of an isotype-matched rhesus recombinant IgG mAb. All animals had been infected with LASIV for different lengths of time, as they were used in previous studies of T cell mediated control of acute SIV replication (42). Four weeks after the CD8β255R1 or IgG control antibody was administered, animals were challenged intravenously with 100TCID50 (0.71ng p27) SIVmac239. Animals were evaluated for an additional 14 weeks before euthanasia, with the exception of animal cy0690, which was sacrificed per veterinarian recommendation at 6 weeks post CD8β-depletion due to non-SIV-related complications.

Flow cytometry was used to monitor CD8αβ+ T cells in PBMC and peripheral LNs of all eleven MCMs. Percent reductions in the absolute count of peripheral CD8αβ+ T cells in animals that received CD8β255R1 were most prominent at week 2 (median: 94%; range: 70% to 98%); CD8αβ+ T cells remained significantly depleted at the time of SIVmac239 challenge (median: 89%; range: 45% to 97%) through the time of necropsy (Figure 2a, blue) (Supplementary Table 1). In control IgG-treated animals, the absolute counts of peripheral CD8αβ+ T cells fluctuated around baseline levels for 16 weeks (Figure 2a, black). Shortly after CD8β255R1 infusion, the reduction in peripheral CD8αβ+ T cell count was statistically significant compared to changes in the IgG control animals (data not shown). Remarkably, the absolute count of peripheral CD8αβ+ T cells at 16 weeks remained depleted by an average of 64% when compared to baseline (median: 67%; range: 28% to 82%) in animals that received CD8β255R1. Persistent depletion of CD8αβ+ T cells after administration of CD8β255R1 lies in stark contrast to the reemergence of CD8α+ lymphocytes within 2 to 4 weeks after administration of the anti-CD8α mAb cM-T807 that has been reported in other studies (8, 10, 14, 31, 40, 45–48, 53–54). Our results in both SIV-naïve and LASIV+ animals are, however, consistent with another report that observed sustained CD8αβ+ T cell depletion in two SIV+ rhesus macaques that were treated with CD8β255R1 and followed for approximately 18 weeks (40).

The extent to which CD8β255R1 depletes LN CD8αβ+ T cells in macaques during either chronic LASIV or pathogenic SIV infection has not yet been defined (40, 41). While technical limitations prevented the longitudinal analysis of cell populations in lymph nodes of individual LASIV+ animals, we compared the percentage of CD8αβ+ T cells in the lymph nodes of animals treated with CD8β255R1 to those treated with the control IgG antibody (Figure 2b). At week 3 post infusion, there were threefold fewer (p<0.006) CD8αβ+ T cells as a proportion of CD3+ T cells in biopsies of axillary lymph nodes isolated from animals infused with CD8β255R1 (median: 4%; range: 1.9% to 20%) compared to control IgG animals (median: 27%; range: 21% to 30%). When lymph nodes were assessed 17 to 18 weeks after mAb infusion following necropsy, CD8αβ+ T cell frequencies remained substantially lower in the majority of CD8β255R1-treated animals compared to control IgG-treated animals (Figure 2c).

### Plasma viral load increases transiently following CD8β255R1 infusion

To determine the impact of CD8β depletion on control of chronic LASIV viremia we measured levels of circulating virus after mAb infusion. Plasma viral load peaked 3 to 11 days after infusion with CD8β255R1 and was significantly increased at day 7 (p<0.001), day 10 (p<0.001), and day 14 (p=0.006) compared to baseline (Figure 3a). Surprisingly, at weeks 3 and 4 post infusion with CD8β255R1, viral loads returned to levels that were not significantly different from baseline, despite diminished CD8αβ+ T cells in peripheral blood and LNs (Figure 2). As expected, minimal changes to peripheral CD8αβ+ T cells during the first four weeks after administration of the control IgG antibody (Figure 2) corresponded to viral loads that remained essentially unchanged (Figure 3b).

### Minimal changes to CD4+ T cells following infusion with CD8β255R1

One previously reported consequence of depletion of CD8α+ cells was a corresponding increase in the frequency or activation state of CD4+ T cell populations (12, 45–47). We monitored the frequency and proliferation of peripheral CD4+ naive (T_N_), CD4+ effector memory (T_EM_), and CD4+ central memory (T_CM_) T cells following CD8β255R1 infusion. Changes from baseline for both the absolute count (Figure 4a) and percent Ki-67+ cells (Figure 4b) were compared to baseline within and between each antibody-treated group. Although peripheral CD4+ T_EM_ cells were significantly increased compared to baseline at two time points in the CD8β255R1-treated group (Figure 4a, middle), these changes did not differ significantly when compared to animals that received control IgG (data not shown). Similarly, CD4+ T cell subpopulations exhibited minimal changes in proliferation compared to baseline within and across groups (Figure 4b and data not shown). We also compared changes in the frequency of total (Figure 4c) and percent Ki-67+ (Figure 4d) CD4+ T cells in the lymph nodes isolated at 3 weeks from animals in both groups. We found that the frequency of CD4+ T cells was higher (p=0.029; Figure 4c) in the CD8β255R1-treated animals compared to control IgG animals, but that may be attributed to a lower frequency of lymph node CD8αβ+ T cells detected at this time (Figure 2b). The percent Ki-67+ of total CD4+ T cells was similar between groups at week 3 post mAb infusion (Figure 4d) (Supplementary Table 2). From these analyses, animals who received an infusion of CD8β255R1 exhibited minimal changes in the number and proliferative capacity of CD4+ T cells, even while plasma viremia was fluctuating.

### Proliferation of CD8αα+ T cells, NK cells, and γδ T cells after infusion with CD8β255R1

Another consequence of using a CD8α-specific mAb is the simultaneous depletion of additional cell populations that express CD8α, such as CD8αα+ T cells, NK cells, and γδ T cells (10, 12, 14, 31, 40, 47, 53). Following infusion with CD8β255R1, we evaluated the absolute count and percent Ki-67+ of these three cell populations in the periphery. In animals that received CD8β255R1, there was a rapid increase in the absolute count of CD8αα+ T cells (median: 3.6-fold; range: 3.2-fold to 3.8-fold) that differed initially from control IgG animals (Figure 5a, top) (Supplementary Table 3). This was followed by increases in the percent of CD8αα+ T cells expressing Ki-67 (median: 4.5-fold; range: 4.2-fold to 5.3-fold) (Figure 5a, bottom). Control IgG-treated animals did not exhibit significant differences from baseline in the absolute count of CD8αα+ T cells or percent that express Ki-67. Few differences were detected in the absolute count of NK cells (Figure 5b, top) or γδ T cells (Figure 5c, top) in each group or between groups for four weeks after infusion (Supplemental Table 3). In contrast, we detected significant increases from baseline in the frequency of NK cells and γδ T cells expressing Ki-67 in the group treated with CD8β255R1 that peaked at approximately 7-fold (median: 4.3-fold; range: 1.8-fold to 17.7-fold) and 10-fold (median: 9.9-fold; range: 0.1-fold to 18.8-fold), respectively (Figures 5b and 5c, bottom). The observed proliferation of NK cells in the CD8β255R1-treated animals was significantly greater than control IgG-treated animals at days 7, 10, and 14 (Supplementary Table 4).

We also evaluated changes to the frequency and proliferation of CD8αα+ T cells (Figure 5d), NK cells (Figure 5e), and γδ T cells (Figure 5f) in lymph nodes of animals treated with CD8β255R1 or control IgG at week 3 post infusion (Supplementary Table 2). Similar to cells in the periphery, the frequency of CD8αα+ T cells (Figure 5d, top), NK cells (Figure 5e, top), and γδ T cells (Figure 5f, top) in lymph nodes of animals treated with CD8β255R1 was comparable to control IgG animals. At week 3 after CD8β255R1 infusion, the frequency of Ki67+ CD8αα+ T cells (Figure 5d, bottom) and γδ T cells (Figure 5f, bottom) was significantly higher than that observed in control IgG animals. Taken together, our data indicate that CD8αα+ T cells, NK cells, and γδ T cells exhibited rapid proliferation in either the PBMC or lymph nodes following CD8β255R1 infusion, while their absolute cell counts remained relatively unchanged.

### Limited SIV-specific T cell responses in LNs following CD8β255R1 infusion

To determine the impact of CD8β255R1 infusion on SIV-specific T cells within LNs, we performed intracellular cytokine staining (ICS) with cells isolated from LN biopsies taken three weeks after antibody infusion. Multiple pools containing overlapping peptides spanning the entire Gag protein and Env protein were used to stimulate cells, followed by staining with CD107a (Figure 6, top) and IFNγ/TNFα (Figure 6, bottom) to assess degranulation and pro-inflammatory cytokine secretion, respectively. None of the animals that received the control IgG antibody exhibited remarkable SIV-specific responses. This was not surprising, as these animals did not exhibit any marked changes in viremia. We were surprised to find that three of the animals that received CD8β255R1 (cy0691, cy0749, and cy0752) exhibited more residual virus-specific CD8αβ+ T cells than the others (Figure 6). Of note, those three animals were each infected with a LASIV that contained known immunogenic epitopes. Thus, ongoing replication of a virus with intact M3 epitopes may have provided stronger antigenic stimulation to T cells and therefore driven proliferation in these three animals. The other four animals that received CD8β255R1 were homozygous for the M3 MHC haplotype and originally infected with a LASIV that contained point mutations in eight epitopes restricted by MHC molecules expressed by the M3 MHC haplotype (42). Even though it is likely that CD8β255R1 infusion alone cannot completely eliminate virus-specific CD8αβ+ T cells from lymph nodes, the combination of infecting animals with a minimally immunogenic LASIV and infusion of CD8β255R1 was able to limit the frequency of Gag- and Env-specific CD8β+ T cells at this important tissue site.

**Figure 6.**
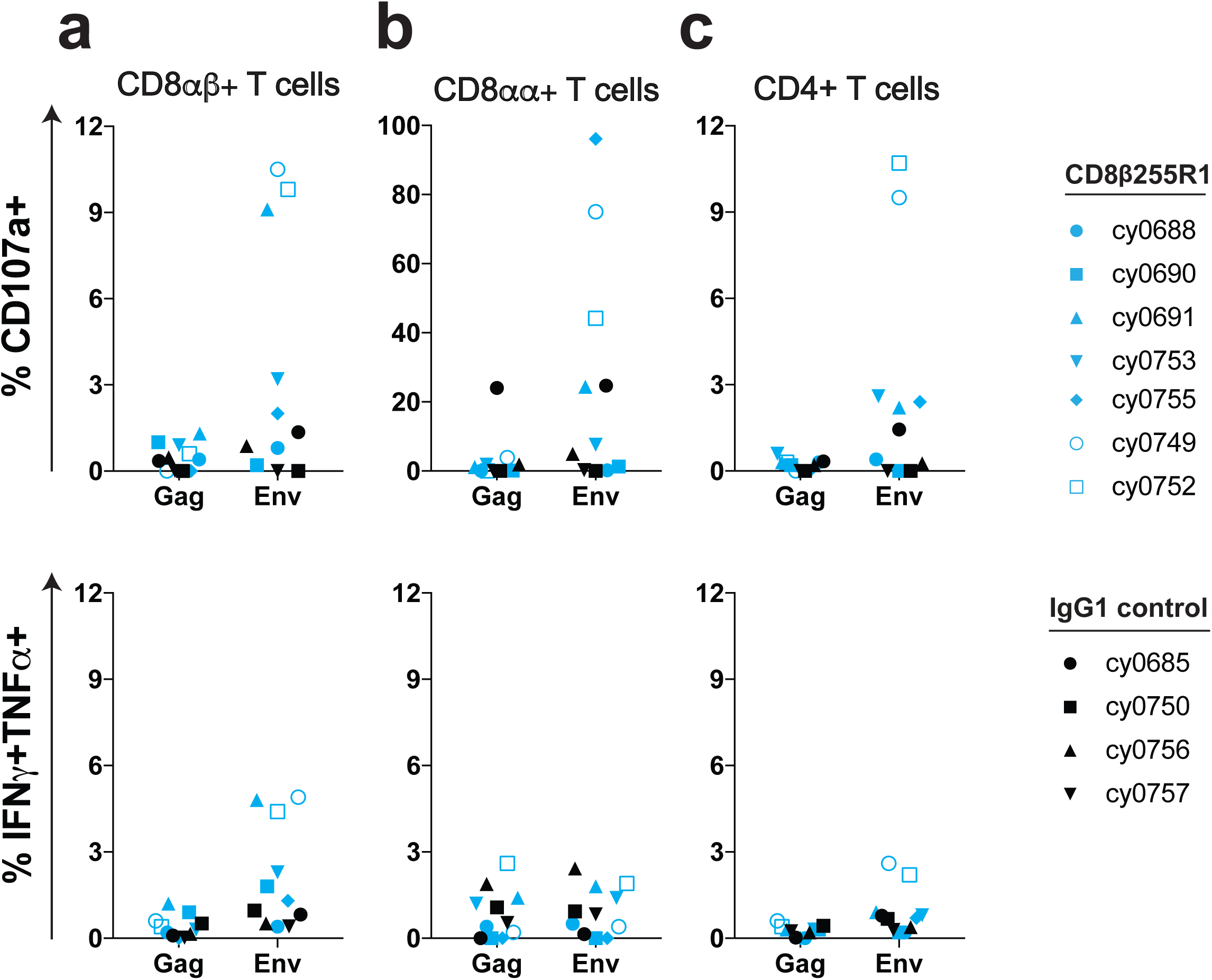
Env-specific antiviral responses in lymph nodes following infusion with CD8β255R1. At 3 weeks post-infusion we performed intracellular cytokine staining to assess Gag-specific and Env-specific responses from lymph node biopsies for (a) CD8αβ+ T cells, (b) CD8αα+ T cells, and (c) CD4+ T cells. We assessed degranulation (CD107a, top) and pro-inflammatory cytokine secretion (IFNγ and TNFα, bottom) for seven animals infused with CD8b255R1 (blue) and four animals infused with IgG1 control (black). Both single and double positive responses were included as % IFNγ+TNFα+. Symbols represent individual animals.

### Protection from SIVmac239 challenge during depletion of CD8αβ+ T cells

To determine whether CD8β255R1 infusion increased susceptibility to infection with pathogenic SIV, we performed a high dose intravenous challenge in all 11 animals at 4 weeks post antibody infusion with SIVmac239 (55, 56). Previously, we found that 4/7 MCMs chronically infected with a minimally immunogenic LASIV were protected from intravenous challenge with the same dose of SIVmac239 (34). The 11 animals used in the current study were also chronically infected with a *nef*-deleted SIV, so discriminating qRT-PCR assays were used to differentiate previous Δ*nef* circulating virus from full-length *nef* that was present in the challenge strain. Surprisingly, there was no replicating SIVmac239 in the plasma of six of the seven MCMs infused with CD8β255R1 (Figure 7a), despite depletion of peripheral CD8β+ lymphocytes at the time of challenge (median: 87%; range: 45% to 95%) (Figure 2a). Similarly, there was no replicating SIVmac239 detected in the plasma of three out of four MCMs that received the control IgG antibody (Figure 7b), despite having similar CD8αβ+ T cell levels as pre-depletion (Figure 2a). It is possible that residual virus-specific CD8αβ+ T cells in the lymph nodes (Figure 6) were sufficient to protect animals from challenge, as others have found that virus-specific CD8+ T cells in lymph nodes are an immune correlate of protection by LASIV (38). However, with so few animals becoming infected with SIVmac239 we could not determine if this was the main mechanism of protection.

**Figure 7.**
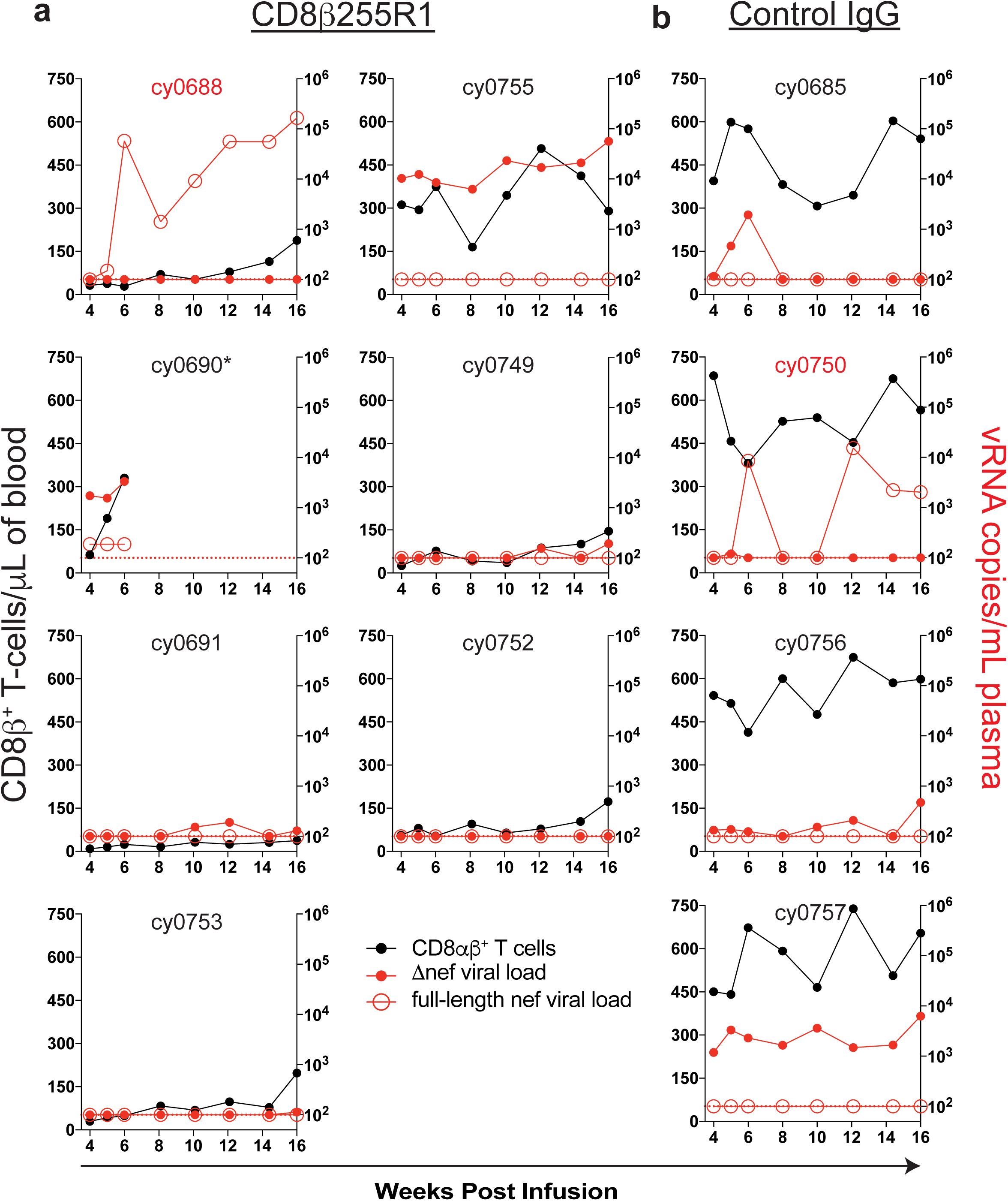
Protection from intravenous SIVmac239 challenge following infusion with CD8β255R1. Four weeks after infusion with CD8β255R1 or control IgG, animals were challenged intravenously with SIVmac239. Viral load assays were performed for full-length *nef* (red, closed) and Δ*nef* (red, open) at the indicated time points for each animal and graphed in the context of absolute CD8αβ+ T cell count (black). The limit of detection for the assay (100 vRNA copies/mL) is represented by a horizontal red dotted line. Animal IDs highlighted in red became infected following SIVmac239 challenge. *cy0690 required necropsy at week 6 following CD8β255R1 infusion.

### Neutralizing antibody titers to SIVmac239 do not predict challenge outcome

We wanted to determine whether depletion of CD8β+ lymphocytes had a direct impact on neutralizing antibody titers, which may have contributed to protection from pathogenic SIVmac239 infection. To accomplish this, we performed *in vitro* antibody neutralization assays with plasma collected immediately before mAb infusion, as well as immediately before SIVmac239 challenge. We tested neutralization potency against SIV strains that are highly neutralization-sensitive (tier 1; SIVmac251.30 and SIVsmE660.CP3C), moderately neutralization-resistant (tier 2; SIVmac251.H9), and highly neutralization-resistant (tier 3; SIVmac239) (43). In animals that received control IgG, neutralizing antibody titers prior to infusion (Figure 8a, left) were similar to those detected prior to SIVmac239 challenge (Figure 8a, right). Similarly, we did not detect substantial changes to neutralizing antibody titers prior to CD8β255R1 infusion (Figure 8b, left) when compared to the titers right before SIVmac239 challenge (Figure 8b, right). Accordingly, we could not conclude protection from challenge following infusion with CD8β255R1 was a result of increased *in vitro* SIVmac239 neutralization antibody titers.

**Figure 8.**
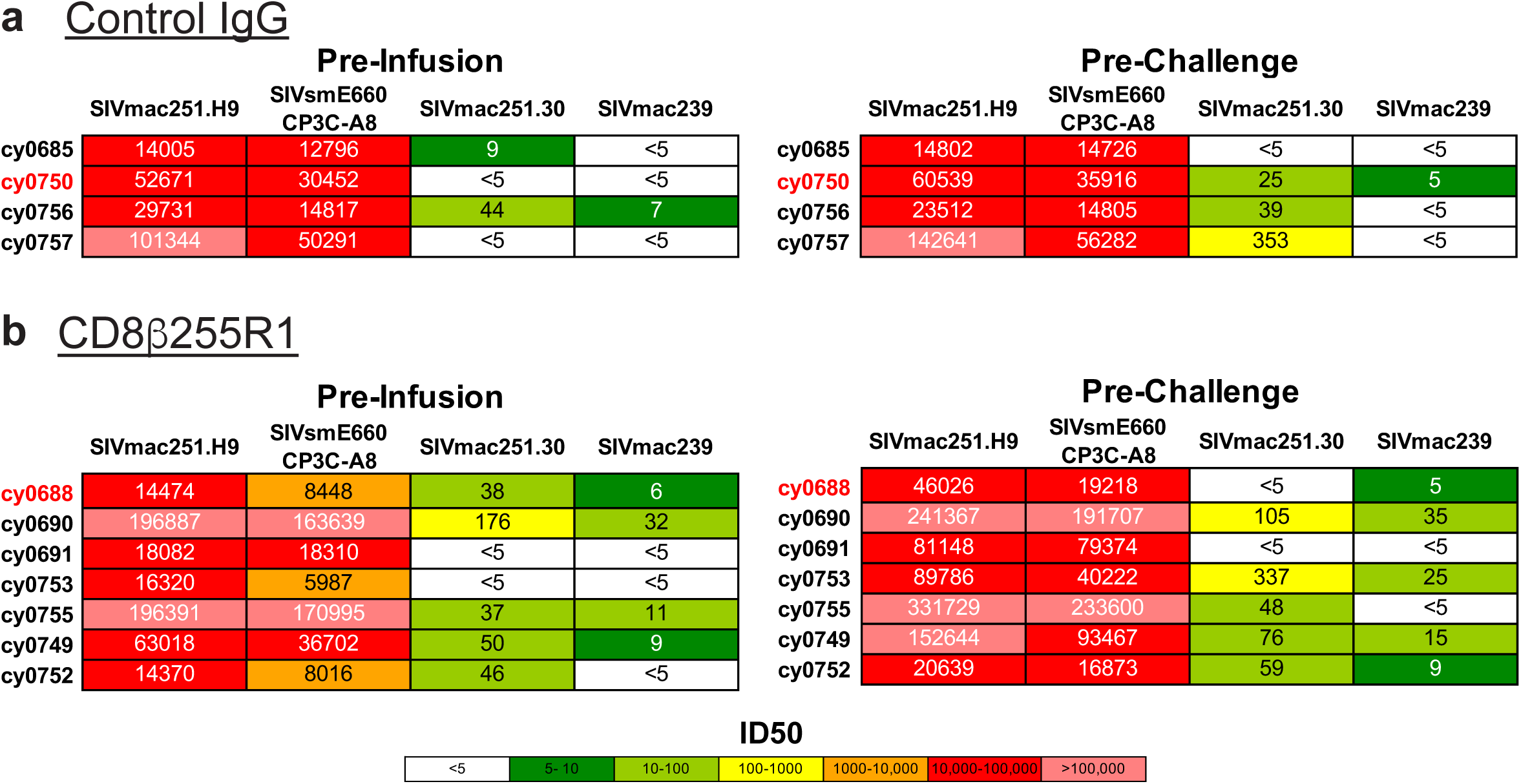
Plasma neutralization potency *in vitro* is not predictive of protection from challenge *in vivo*. Plasma samples collected immediately before mAb infusion and 4 weeks later immediately before intravenous SIVmac239 challenge were used to test antibody neutralization potency against select tier 1 (SIVmac239.H9, SIVsmE660.CP3C), tier 2 (SIVmac251.30), and tier 3 (SIVmac239) viruses. Neutralizing antibody titers were measured (a) prior to infusion with control IgG and (b) SIVmac239 challenge. Neutralizing antibody titers were measured (c) prior to infusion with CD8β255R1 and (d) SIVmac239 challenge. Titers are shown as 50% inhibitory dilution (ID50). Animals that became infected following SIVmac239 challenge are highlighted in red.

## Discussion

Studies of SIV-infected macaques using a CD8α-specific mAb to deplete CD8+ T cells *in vivo* have frequently been interpreted as evidence for a direct antiviral role of CD8+ T cells in HIV infection (8, 48). However, these results are complicated by the simultaneous depletion of CD8α-expressing NK cells and γδ T cells (10, 14, 15, 49). Here we showed that a single intravenous infusion of the CD8β-specific mAb CD8β255R1 selectively depleted CD8αβ+ T cells in peripheral blood, though to a lesser extent in lymph nodes, from SIV-naïve and LASIV-infected cynomolgus macaques. Following depletion, we observed a transient rise in plasma viremia that was followed by re-establishment of viral control, even when CD8αβ+ T cells remained depleted in the peripheral blood. During the time of prolonged peripheral CD8αβ+ T cell depletion, we found that CD8αα+ T cells, NK cells, and γδ T cells exhibited markers of proliferation regardless of changes to their total frequencies in blood. These observations demonstrated that the CD8β255R1 antibody specifically depleted CD8αβ+ T cells. CD8β255R1 could not fully deplete CD8αβ+ T cells from tissue sites, as lymph nodes contained residual CD8αβ+ T cells. While our results specifically demonstrated that administration of the CD8β255R1 depleting antibody led to a rise in LASIV plasma viremia, we could not conclusively determine whether the residual CD8αβ+ T cells in tissues, or the persistence of CD8αα+ T cells, NK cells, and/or γδ T cells were responsible for re-establishment of LASIV control and subsequent protection from pathogenic SIVmac239 challenge. Because animals also had very low neutralizing antibody titers against SIVmac239 throughout the study period, it is unlikely that protection against challenge was mediated by neutralization of the challenge virus.

We found that animals treated with the CD8β255R1 antibody exhibited demonstrably fewer total CD8αβ+ T cells in the periphery and the lymph nodes, when compared to control IgG animals. To determine if we depleted virus-specific CD8αβ+ T cells, we initially performed γ-interferon enzyme-linked immunospot assays (IFN-γ ELISPOT) with PBMC prior to, and three weeks after infusion with CD8β255R1, but were unable to confirm if the SIV-specific responses that were detected three weeks after infusion were attributed to CD8+ T cells or CD4+ T cells (data not shown). While numerous pieces of evidence point towards a critical role of CD8+ T cells in long term viral control (8–10, 14, 31, 47, 53), one report suggests that virus-specific CD8+ T cells in peripheral blood may not be absolutely required for control of virus replication during chronic SIV infection, even if they are needed to initially suppress viremia (57). While tetrameric reagents are an alternative method to detect virus-specific T cells, these reagents were not available for two reasons: (1) Many of the epitopes that would elicit CD8αβ+ T cells were rendered non-immunogenic in the mutant LASIV used to infect several of these animals (42), and (2) tetrameric reagents specific for the CD8 and CD4 T cells that developed in the animals infected with the mutant LASIV have not been produced. We did perform intracellular cytokine staining experiments with lymph nodes that were collected three weeks after antibody infusion. Interestingly, the three animals with the largest frequency of Env-specific CD8αβ+ T cells were the three animals that were originally infected with strains of LASIV containing immunogenic epitopes. Two animals were infected with wild type SIVmac239Δnef, and the third animal was infected with the mutant LASIV, but expressed some non-M3 MHC alleles. As a result, these three animals likely had the largest pool of virus-specific memory CD8αβ+ T cells that could emerge when viremia increased. Together, our data imply that the most effective way to eliminate virus-specific CD8αβ+ T cells with currently available technology is to infect animals with a virus whose T cell epitopes are rendered non-immunogenic combined with a CD8β-specific depleting antibody. Even with this ‘double-knockout approach’, more sophisticated reagents are needed to improve CD8αβ+ T cell depletion in tissues to determine if they are required to re-establish viral control.

Even when we minimized the CD8αβ+ T cell populations with our interventions, we were surprised to find that six out of seven animals treated with CD8β255R1 were resistant to infection with pathogenic SIVmac239. This level of protection was on par with the animals treated with control IgG. Unfortunately, even when we examined non-CD8β+ immune cell populations and neutralizing antibody titers, there were no obvious immune correlates of protection. Protection afforded to LASIV has previously been associated with the presence of effector-differentiated T cell responses within the lymph node (37, 38), so it is entirely possible that the residual virus-specific T cells in the lymph nodes or tissues were sufficient to protect animals from SIVmac239 challenge. Continuing to improve the methods to deplete virus-specific immune cells in the LASIV model will be needed to directly demonstrate the immune correlates of protection mediated by LASIV.

One unique attribute of our study was the inclusion of animals treated with an isotype-matched IgG control antibody that was absent in recent reports of CD8β255R1 (40, 41). By comparing paired differences of cell populations between animals treated with CD8β255R1 and control IgG, we accounted for potential artifacts that may result from infusion of a nonspecific IgG antibody. For example, increases over baseline in the percent of Ki-67+ CD8αα+ T cells detected from weeks 2 to 4 in both groups suggests that IgG infusion alone induces proliferation of CD8αα+ T cells. Additionally, similar to a previous study using CD8β255R1 (40) we observed increased numbers of circulating NK cells in animals receiving CD8β255R1, though similar increases were also detected in the control IgG animals. These observations question whether changes to CD8αα+ cell populations were a direct consequence of administration of the specific CD8β255R1 monoclonal antibody, or whether the administration of a control antibody was sufficient to induce this expansion. Thus, including an IgG control group in these types of antibody-infusion studies is critical for identifying non-specific depletion effects.

In this study, we provide a comprehensive comparison of the immunological impact that follows infusion with the anti-CD8β mAb CD8β255R1 compared to an isotype matched control IgG. The persistent depletion of peripheral CD8αβ+ T cells following administration of CD8β255R1 lies in stark contrast to the transient depletion of CD8αβ+ T cells following infusion with a CD8α-specific mAb. Unlike studies with the CD8α-depleting antibody that observed regained control of virus replication coincident with rebound of CD8+ T cells, we observed regained control of viral replication, even when peripheral CD8β+ T cells remained depleted. Moreover, protection from intravenous challenge with pathogenic SIVmac239 was achieved when CD8αβ+ T cell magnitude was reduced by administration of CD8β255R1 and, in some cases, in combination with a mutant LASIV that failed to elicit many virus-specific T cells (42). Nonetheless, the data we provide demonstrates that the CD8β255R1 antibody can be used to specifically deplete CD8αβ+ T cells, while leaving other CD8α+ immune cell populations intact. This may serve to be valuable in future studies evaluating the importance of CD8αβ+ T cells in diverse disease models.

## Acknowledgements

Research reported in this publication was supported in part by the Office of The Director, National Institutes of Health (NIH), under award number P51OD011106 to the Wisconsin National Primate Research Center (University of Wisconsin-Madison), as well as by the Intramural Research Program of the National Institutes of Allergy and Infectious Diseases, NIH. The research was conducted in part at a facility constructed with support from Research Facilities Improvement Program grant numbers RR15459-01 and RR020141-01. We would like to thank staff at the Wisconsin National Primate Research Center (WNPRC) for veterinary care of the animals involved in the study. Research reported in this publication was supported in part by National Institutes of Health award number R01AI108415, as well as by the National Institute of General Medical Sciences of the National Institutes of Health under award number T32GM081061. The content is solely our responsibility and does not necessarily represent the official views of the National Institutes of Health. Special thanks to Keith Reimann, Afam Okoye, Mauricio Martins, and Matthew Reynolds for helpful discussions.

